# The effect of extended cold storage and the use of extenders on motility and swimming kinematics of shortnose sturgeon sperm

**DOI:** 10.1101/2024.08.18.608476

**Authors:** Kenzie H. Melanson, Matthew K. Litvak

## Abstract

Shortnose sturgeon are listed as a species of special concern in Canada and as endangered in the US. Increasing knowledge about this species, particularly in the area of reproductive biology, will better the management of wild populations and aid in the development of assisted reproduction protocols. However, access to wild sperm is limited, so short-term and long-term storage of sperm from sturgeon is crucial for reproductive studies. Here we report on testing and development of a short-term storage protocol for shortnose sturgeon. Milt samples were collected from wild shortnose sturgeon caught in the Wolastoq River. Subsets of semen were mixed with different extenders with or without oxygen; control treatments without extenders were also run. We used computer-assisted sperm analysis (CASA) to determine sperm motility and swimming kinematics for the different treatments. All groups were examined immediately after collection and treatment application, and then 1, 2, and 7 days after storage in a fridge (4°C) for experiment 1, and days 1, 3, 7, 10, 14, 17, 21, and 24 for experiment 2. The response variables motility, curvilinear velocity (VCL), linearity (LIN), and wobble (WOB) showed an overall decrease over time with differences between extender treatments. While untreated milt maintained some motility up to day 21, the addition of an extender reduced decline in motility and improved longevity up to day 24. Milt treated with the Park and Chapman extender had the slowest motility decline of extenders used, and milt treated with the modified Tsvetkova extender showed less potential for contamination.

## 1. Introduction

Sturgeon, valued for their caviar and meat, are the most endangered vertebrate group in the world, with all 27 species listed on the International Union for Conservation of Nature (IUCN) Red List (IUCN, 2023). This is due to habitat fragmentation from dams, water regulation, pollution, and fisheries persecution, through either directed fisheries and/or by-catch or poaching (Litvak, 2010; Bronzi et al., 2011).

Shortnose sturgeon (*Acipenser brevirostrum*) has been listed as a species of special concern since 1980 and endangered in the US since 1967 (COSEWIC, 2015). Shortnose sturgeon in Canada and the US are protected from commercial harvesting, however they are still vulnerable to threats that affect other sturgeon species (DFO, 2014; COSEWIC, 2015; Kynard et al., 2016). Declines in sturgeon have increased interest in aquaculture development and conservation (Giberson and Litvak, 2003; Litvak, 2010; Bronzi et al., 2011).

In many endangered fish species, including sturgeon, milt quantity is limited in availability (Gilroy and Litvak, 2019) due to low numbers and a short breeding season. This is exacerbated by lack of synchronization of egg and sperm production in both the lab and the field settings (Kime et al., 2001; Butts et al., 2012). Development of protocols for both short-term and long-term storage of sperm are crucial for artificial reproductive efforts in conservation and aquaculture (Alavi and Cosson, 2005; Barulin and Shumski, 2022). Short-term storage of refrigerated milt is inexpensive. Stored milt can be used for assisted reproduction, sperm quality analysis, and development of cryopreservation protocols (Contreras et al., 2020). This is particularly important to defeat timing availability issues of sperm and eggs and for control of aquaculture and conservation genetic diversity. However, few short-term sperm storage protocols exist for fishes.

Sturgeon sperm, like that of teleost fish, are immotile in the testis and in seminal plasma (Ginzburg, 1972; Alavi and Cosson, 2005). Activation is controlled by hypo-osmolality or a decrease of extracellular K+ ions when sperm are exposed to water during reproduction (Gallis, 1991; Alavi et al., 2004; Alavi et al., 2012; Alavi et al., 2019). Several factors can affect the quality and viability of stored sperm, such as individual male variability and storage conditions (Aramli et al., 2013). Other factors include temperature, sperm concentration, and oxidative stress (Barulin and Shumski, 2022), as well as types of extenders (Park and Chapman, 2005).

Extender solutions serve as diluents to increase the volume of the samples, protect the sperm cells, and provide optimal conditions as a source of energy for fertilization (Park and Chapman, 2005). Extenders should mimic the seminal fluid composition, as ions at the right concentration will prevent motility activation and maintain osmotic equilibrium between the medium and the sperm cells (Bobe and Labbe, 2008; Alavi et al., 2019). Park and Chapman (2005) developed an artificial extender to be used with milt from Gulf of Mexico sturgeon and shortnose sturgeon. Other extenders for sturgeon semen include Hank’s Balanced Salt Solution (HBSS) (Park and Chapman, 2005), and modified Tsvetkova (Dorsey et al., 2011), as well as other combinations of ions at similar osmolalities (Shazada et al., 2024).

Here, we focused on comparing the effect of different extenders and oxygen on the motility and swimming kinematics of shortnose sturgeon sperm over several days. We ran two experiments. Experiment 1 (2022) compared motility and swimming kinematics of control samples (no extenders) with those treated with the Park and Chapman extender or HBSS extenders. Experiment 2 (2023) compared sperm motility and swimming kinematics of control samples (no extenders) with those treated with the Park and Chapman extender or the modified Tsvetkova extender, and all treatments with and without pure oxygen.

## 2. Materials and methods

### 2.1. Animal care protocol

This study was performed under the guidelines of the Canadian Council of Animal Care: Mount Allison Animal Care Protocols 102193 and 103376.

### 2.2. Fishing

Shortnose sturgeon were caught using gillnets (DFO Scientific Fishing Permit 330697) with 5” stretch mesh between May 13^th^ and May 20^th^, 2022 for experiment 1, and May 19^th^ and 20^th^ 2023 for experiment 2, in the lower reaches of the Wolastoq River (aka Saint John River) in New Brunswick, 11 km below the Mactaquac dam. Fishing was attempted three times for experiment 1 (21 fish total), and four times (18 fish total) for experiment 2, with nets being left in the river between 30 and 60 minutes per attempt. Individuals were placed in a 400 L circular holding tank on the boat and transferred two 1035 L insulated tanks (D337 Saeplast) onshore. A pump supplied the onshore tanks with river water. All fish were tagged using passive integrated transponder (PIT: Biomark) tags for identification during sperm collection. Five males were used for experiment 1 and four males were used in experiment 2.

### 2.3. Sperm collection

Fish were taken out of the holding tank and identified with a PIT tag reader (Biomark HPR Lite). Most fish were already spermiating during handling, so no other identification was necessary to determine the sex of the fish. Shop towels were used to gently dry the urogenital opening to avoid urine contamination. Collection syringes (either 20 or 60 ml) were equipped with a ∼10 cm length of 1/8” inner diameter, 1/4” outer diameter Tygon tubing to collect sperm. A small cut was made near the end of the tubing to prevent the tubing from suctioning to the urogenital canal. If there was not an immediate inflow of milt when the syringe was drawn back, slight pressure was applied to the abdomen to stimulate the flow of milt. For all fish, milt was continuously collected until the syringe could not be drawn back any further. Once collected, syringes with milt were taken to a 20×8’ mobile lab, equipped with power, lights, and an air conditioner, and immediately put into a fridge (∼4°C). Analysis was conducted immediately after milt collection. Milt was collected from fish one at a time, to ensure that the application of treatments and analysis would be done immediately.

### 2.4. Treatments

#### 2.4.1. Experiment 1

The extenders used for experiment 1 were HBSS (1X 21- 020-CM, diluted to ∼100mOsm) (Corning, 2013) and the Park and Chapman extender (Park and Chapman, 2005). There were six treatments in total per fish (Figure 1). Tygon tubing was used to attach the collection syringe to a new 20ml syringe, and 1/6 of collected milt was dispersed into the new syringe, Tygon tubing was reused so long as there was only milt from the same fish being transferred. Milt was kept in the fridge until it was needed. Four 50 ml centrifuge tubes were filled with one of two extenders at a 1:2 (milt to extender) ratio to the measured milt in the syringes. Four of six syringes were individually added to one of four tubes, so each tube had 1/6 amount of milt and an extender. These tubes were shaken to mix, and then pulled back into the labeled syringes. The two syringes with no extender had the same amount of milt but were left undiluted. Each syringe contained less than 10 ml of final solution to ensure each syringe was half empty to allow air exchange and circulation. Syringes were stored in Ziploc bags in the fridge, with the plunger of the syringe towards the coldest part of the fridge. Air exchange was done twice a day for each syringe, by pushing out the old air and sucking in new air. This was done in the morning and evening. Samples were only taken out of the fridge when analysis was being conducted, and they were immediately placed back in the fridge once aliquots were taken.

**Figure 1.**
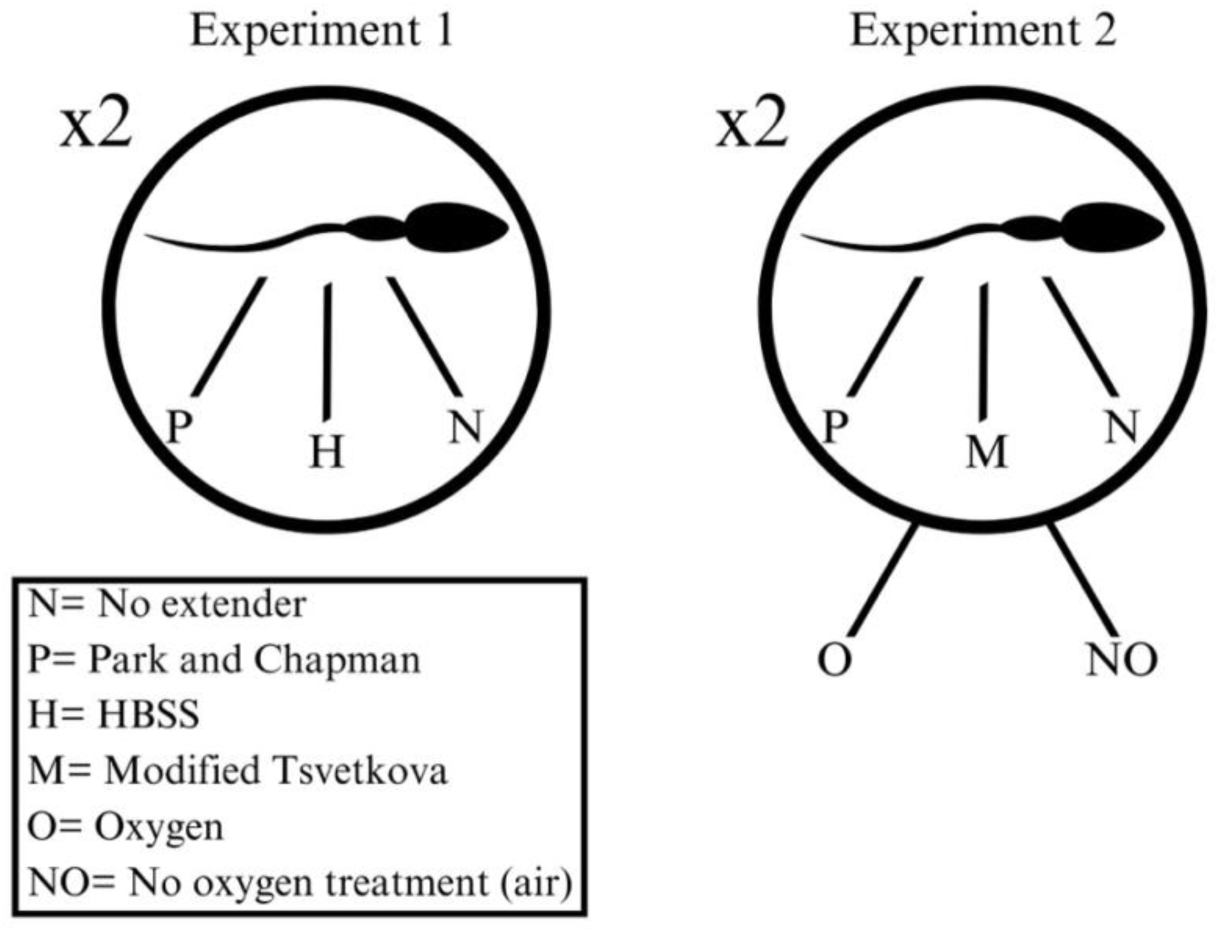
Treatments for experiment 1 and experiment 2. Experiment 1 had six total treatments, experiment 2 had twelve total treatments.

### 2.4.2. Experiment 2

The extenders used for experiment 2 were modified Tsvetkova (Dorsey et al., 2011: 600 ml of a solution made with tris acid buffer and 40 ml of a solution made with tris base buffer, to achieve a neutral pH) and the Park and Chapman extender (Park and Chapman, 2005). There were twelve treatments in total per fish (Figure 1). Tygon tubing was used to attach the collection syringe to a new 20 ml syringe, and 1/12 of collected milt was dispersed into the new syringe, as in Experiment 1, Tygon tubing was reused so long as there was only milt from the same fish being transferred. Milt was kept in the fridge until it was needed. Eight 50 ml centrifuge tubes were filled with one of two extenders at a 1:3 (milt to extender) ratio to the measured milt in the syringes.

Eight of twelve syringes were individually added to one of eight tubes, so each tube had 1/12 amount of milt and an extender. These tubes were shaken to mix, and then pulled back into the labeled syringes. The four syringes with no extender had the same amount of milt but was left undiluted. Six of twelve syringes were treated with oxygen using Tygon tubes fed into a Ziploc bag full of oxygen. All twelve syringes contained less than 10 ml of final solution to ensure each syringe was half empty to allow air exchange and circulation. All syringes were stored in Ziploc bags in the fridge, with the plunger towards the coldest part of the fridge. Air exchange was done after each analysis (every 3-4 days) by pushing out the old air/oxygen and drawing in new air/oxygen. Samples were only taken out of the fridge when the analysis was being conducted, and they were immediately placed back in the fridge once aliquots were taken for analysis.

### 2.5. Analysis of Sperm Activity

Motility and kinematic parameter analysis of sperm was conducted with an Olympus CX41 microscope and CEROS II (Hamilton Thorne) CASA system at a magnification of 20x using 20 μm Hamilton Thorne Inc. 2X-CEL slides. Sperm density for each fish was calculated by adding 2 μL of undiluted sample and 18 μL of 1% bovine serum albumin (BSA), for activation and to prevent sticking of sperm, to a blank slide. A different dilution was used for each fish, calculated to reach a target visible cell count of 40-60 sperm per field of view. During analysis slides were kept at river temperature (8°C) using a Linkam PE95/T95 system controller. The room temperature and humidity was controlled with an air conditioner and de-humidifier to prevent condensation on the cooled slides. 2.0 mL vials and 0.5 mL vials were labelled with the treatment type and each replicate. These vials were placed in a microtube holder which was placed on a paper towel and plastic sheet over ice in the cooler. The plastic bag and paper towel ensured no direct contact of the holder/vials with ice. A few drops (1-3) of milt were added to each 0.5 mL labeled vial, corresponding to the treatment in the syringe, and the syringes were placed back in the fridge. BSA was then measured into each 2.0 mL vial in correspondence to the values calculated in an Excel sheet to reach the target dilution (as mentioned previously) based on sperm density for each fish. Samples for analysis were mixed one at a time to avoid activation until they could be analyzed. The calculated amount of sample to reach the appropriate dilution was taken from the 0.5 mL vial and added to the 2.0 mL vial. The 2.0 mL vial with sample was then shaken slightly to mix. A 20 μL volume of this mixture was then pipette onto a blank slide and put under the microscope to be analyzed. Video captures of 60 frames/second, lasting ∼2 seconds were taken immediately on the CASA system after a cover slip was placed on, and a stopwatch was started. The sample was followed for 1 minute for both experiments with 3-5 videos captured during this time. These analyses were repeated for each sample on days 0, 1, 2, and 7 for experiment 1, and on days 0, 3, 7, 10, 14, 17, 21, and 24 for experiment 2. A CSV file was created automatically by CASA system at the end of each minute increment, with data from all videos for each minute.

### 2.6. Seminal fluid testing

Seminal fluid testing was performed for both experiments. Milt was spun in a Thermo Scientific Sorvall Legend Micro 17 Microcentrifuge until a sperm pellet formed (∼ 9 minutes on 1000 rpm). Seminal fluid was then separated by pipette into 2 μL tubes. Osmolality was tested using Model 3320 Advanced Instruments Inc. osmometer for both experiments, and pH was tested for experiment 2 only, with a Mettler Toledo pH/ion tester.

### 2.7. Variables analyzed

In addition to motility, we chose to analyze three related swimming kinematic factors: curvilinear velocity (VCL), linearity (LIN), and wobble (WOB). VCL was chosen as it is most representative of the sperm velocity. LIN was chosen as it is the straight-line velocity divided by the VCL. WOB was chosen as it is the average path velocity divided by the VCL. Motility, LIN, and WOB are represented as percentages, and VCL is in μm/s. Sperm were classified as motile if they had a VCL>10 µm/s (Xin et al., 2020; Shazada et al., 2024).

### 2.8. Statistical analysis

CSV files from CASA were combined using a Python script, to create one dataset with all data. Only motile sperm were used to assess kinematics, and motility was calculated as a mean sum of motile and nonmotile sperm. Figures were created using full, untransformed data. Data were analyzed using RStudio (R Core team, 2023), version 4.2.2, and significance was assessed at *P*<0.05. Linear mixed models were made for all four dependent variables (VCL, WOB, LIN and motility), using the function *lmer* in the package *lme4* (Bates et al., 2015). Post-hoc tests to determine differences between extenders were done using *emmeans* (Lenth, 2023), which uses estimated marginal means (Searle et al., 1980) to interpret differences. An ANOVA was run on experiment 2 to examine the interaction between day and extender. Days 14-24 were omitted from experiment 2 linear mixed models due to the large number of zeros, and increased contamination after day 10. An ANOVA was run comparing the last day of motility (VCL<10 µm/s) and post-hoc Tukey tests were run to determine differences. Normality and homogeneity of variances were checked using visual analysis of residuals (histogram, qq-line plot, and plot of points against residuals of the model), and by examining the skewness of the residuals. Dependent variables that had residuals with a skewness that was not symmetric (greater than 0.5 or less than −0.5) or that did not pass visual homogeneity of variance and normality tests were transformed using *BoxCoxTrans* in the package *caret* (Kuhn, 2008). For experiment 1, all dependent variables were transformed except VCL. For experiment 2, WOB, the last day of motility, and day 7 for each variable as well as all days of motility for the experiment 2 ANOVA needed to be transformed.

The percent decrease in motility for each treatment was calculated by dividing the difference in motility from day 0 and day 7 by the starting motility (day 0), and was an average of all fish from each experiment.

## 3. Results

### 3.1. Experiment 1

The number of days sperm were held showed a statistically significant decrease on all parameters (Table 1, Figure 2a). Post-hoc comparisons revealed that VCL, LIN, and WOB on days 0, 1, and 2 were significantly higher than values on day 7. There was a significant effect of extender on VCL, and post-hoc testing revealed that the VCL of sperm treated with either extender decreased at a slower rate than sperm left untreated (Table 1). When comparing the last day of motility between treatments there was no statistically significant effect of extender (Table 2). The percent decrease in motility over the 7-day period was lowest for extender P (41.21%), and highest for no extender (N) (72.38%).

**Figure 2.**
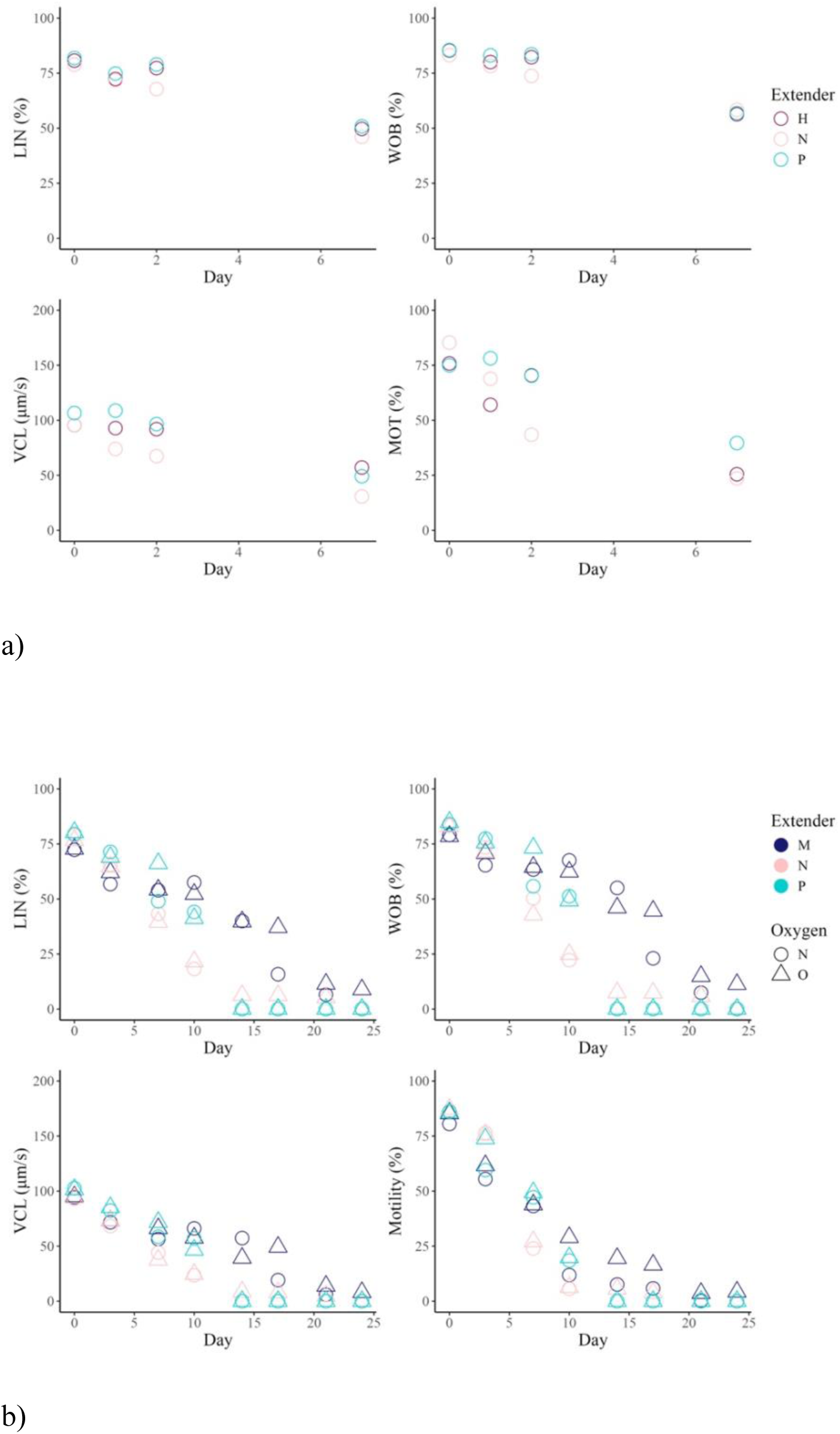
Mean motility, VCL, LIN, and WOB of sperm over time from experiment 1 (a), and experiment 2 (b). Day 0 represents day of sperm collection. Extenders are denoted by colours, H for HBSS, P for Park and Chapman, M for modified Tsvetkova, and N for no extender. Oxygen treatments for experiment 2 are denoted by shapes, O for oxygen treatment and N for no oxygen treatment.

**Table 1.**
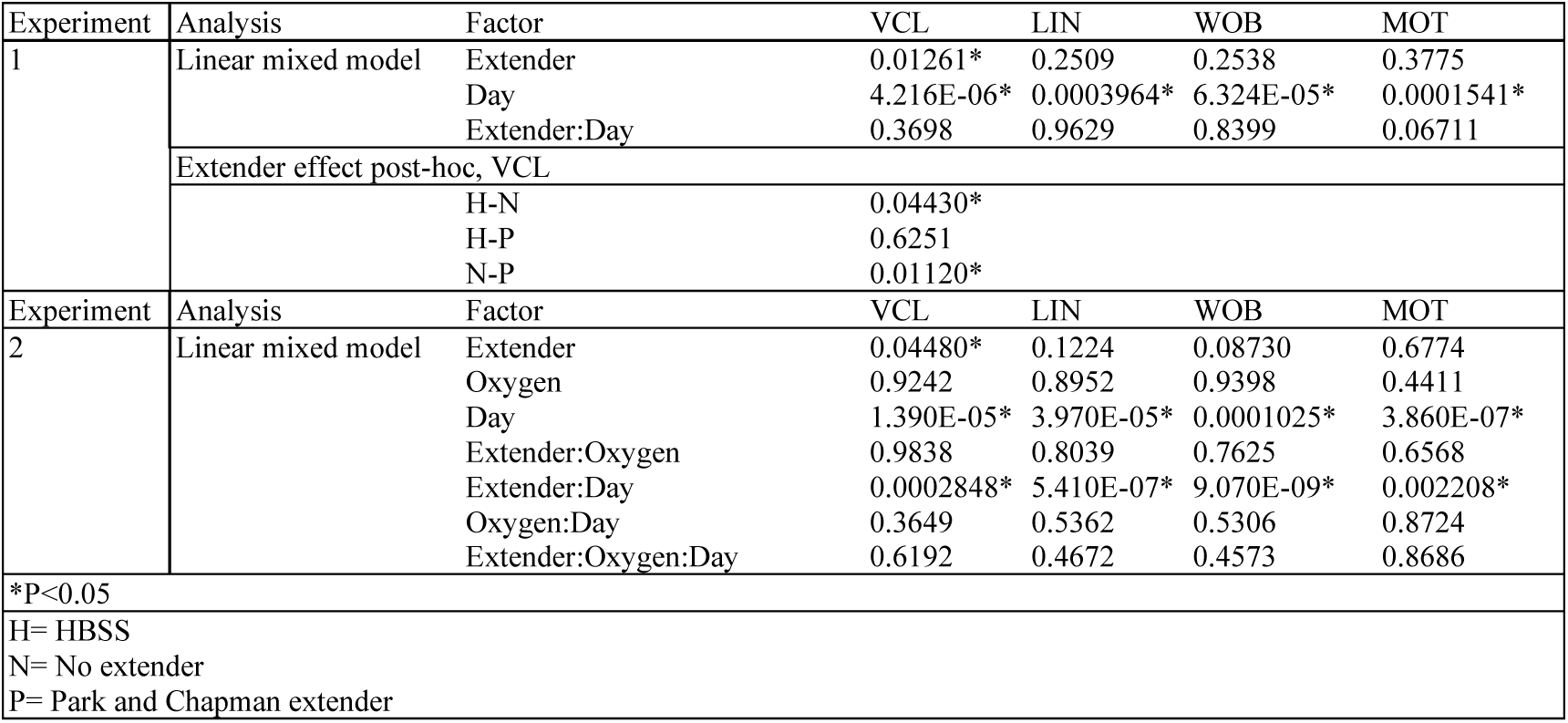
Results from experiment 1 and experiment 2 swimming kinematics statistical analysis using LMER. All parameters decreased over each day. Post-hoc tests are shown for extender effects on VCL for 2022 as there was no interaction.

**Table 2.** Results from experiment 1 and experiment 2 last day of motility ANOVA’s.

| Experiment | Analysis | Factor | Last Day Motile |
| --- | --- | --- | --- |
| 1 | ANOVA | Extender | 0.9350 |
| 2 | ANOVA | Extender | 6.470E-07* |
|  | Tukey | N-M | 1.800E-06* |
|  |  | P-M | 2.270E-05* |
|  |  | P-N | 0.7168 |
| *P<0.05 |  |  |  |
| H= HBSS |  |  |  |
| N= No extender |  |  |  |
| P= Park and Chapman extender |  |  |  |
| M= Modified Tsvetkova extender |  |  |  |

### 3.2. Experiment 2

The amount of days sperm were held showed a statistically significant decrease on all parameters (Table 1, Figure 2b). There were statistically significant interactions between day and extender for all parameters (Table 1) necessitating breaking apart the data and running individual ANOVAs by day. ANOVAs run on each day showed that on day 3 the LIN and WOB of sperm treated with the Park and Chapman extender was significantly higher than that of sperm treated with the modified Tsvetkova extender, and the VCL of sperm treated with the Park and Chapman extender was significantly higher than that of sperm left untreated (Table 3). On day 10, the VCL, LIN, and WOB of sperm treated with either extender was significantly higher than that of sperm left untreated (Table 3).

**Table 3.** Results of Tukey comparisons from the ANOVA for the effect of extenders on each day for experiment 2.

| Day | Comparison | VCL | LIN | WOB | MOT |
| --- | --- | --- | --- | --- | --- |
| 0 | N-M | 0.9447 | 0.07206 | 0.02567* | 0.1814 |
|  | P-M | 0.1322 | 0.008045* | 0.003116* | 0.9869 |
|  | P-N | 0.06773 | 0.6506 | 0.7152 | 0.1356 |
| 3 | N-M | 0.9385 | 0.1590692 | 0.1973831 | 0.8901148 |
|  | P-M | 0.09357 | 0.0026512* | 0.0035696* | 0.3572449 |
|  | P-N | 0.04414* | 0.2272558 | 0.2214556 | 0.1667551 |
| 7 | N-M | 0.3405 | 0.9587 | 0.9137 | 0.9851 |
|  | P-M | 0.6237 | 0.9447 | 0.5914 | 0.9527 |
|  | P-N | 0.8780 | 0.8219 | 0.8326 | 0.9905 |
| 10 | N-M | 0.0002980* | 0.00005600* | 0.00002270* | 0.6938 |
|  | P-M | 0.4979 | 0.2292 | 0.1821 | 0.9878 |
|  | P-N | 0.008559* | 0.008847* | 0.005989* | 0.7825 |
| *P<0.05 |  |  |  |  |  |
| N= No extender |  |  |  |  |  |
| P= Park and Chapman extender |  |  |  |  |  |
| M= Modified Tsvetkova extender |  |  |  |  |  |

When comparing the last day of motility between treatments there was a statistically significant effect of extender, and no statistically significant effect of oxygen. Post-hoc testing demonstrates that motility decreased significantly less for sperm treated with extender P or extender M than for sperm treated with no extender (N) (Table 2). The percent decrease in motility over the first 7 days was lowest for extender P with oxygen (42.42%), and highest for no extender (N) without oxygen (72.29%).

#### 3.2.1. Contamination

Contamination by protozoans was seen on slides as early as day 10, for both oxygen treatments with extender P. All aliquots treated with extender P were contaminated by day 17 regardless of oxygen treatments. Protozoans were seen in one aliquot treated with extender M and no oxygen on day 17, and on day 21 five more aliquots treated with extender M had protozoans visible, four of these were oxygen treated, and by day 24, eight aliquots treated with extender M were recorded as having protozoans. Only three aliquots treated with no extender (N) had visible protozoans, two starting on day 14 and one on day 17. Sperm treated with the Park and Chapman extender had no motility after 14 days, likely due to protozoan contamination. The sperm treated with the modified Tsvetkova extender had minimal protozoan contamination when compared to the Park and Chapman extender.

### 3.3. Osmolality and pH measurements

The modified Tsvetkova extender had a pH of 7.7, and an osmolality of 68. The Park and Chapman extender had a pH of 7.6 and an osmolality of 108 for experiment 2 (only tested in experiment 2). HBSS was adjusted to have an osmolality of around 100. The osmolalities and pH for each extender fell within the range of the osmolalities (40-147 mOsm/kg) and pH (7.95-8.68) for shortnose sturgeon seminal fluid in this study.

## 4. Discussion

The goal of our study was to determine a short-term storage protocol that allows for longevity of sperm motility and swimming kinematics for shortnose sturgeon sperm.

The percent decrease for motility for both experiments indicate that the Park and Chapman extender maintained the highest level of motility and swimming kinematic performance over the first 7 days compared to the other treatments. However, the sperm treated with the modified Tsvetkova extender had better motility and performance from day 10 until the end of the experiment. Sperm treated with either extender performed better than sperm stored without extenders. The last day of motility analysis shows that sperm treated with the modified Tsvetkova extender maintains motility for the longest period of time; it was the only treatment that lasted until day 24. Sperm treated with the Park and Chapman dropped in motility earlier than sperm treated with Tsevtkova, likely due to protozoan contamination. Samples treated with the modified Tsvetkova extender had lower visible protozoan contamination than the samples treated with the Park and Chapman extender or the control non-extender/treated samples. This finding could be important in future extender research as the Tsevtkova extender appears to not only prolong the life of sperm, but also decreases contamination by protozoans.

Even with no treatment, motility did persist up to 24 days, indicating that even without an extender swimming kinematics can be maintained. This is important to note as the main goal of the study was to determine how long sperm can last in short-term storage so it can be transferred to the lab to be cryopreserved, without having to take cryopreservation equipment into the field.

When examining sperm kinematics and parameters, it is important to keep in mind the biological significance of these sperm kinematics in terms of sperm quality. The classification of sperm quality varies by researcher. Motility values less than 20% (Xin et al., 2020) or 30% (Dorsey et al., 2011) have been classified as poor quality, values between 20-80% (Xin et al., 2020) or 30-90% (Dorsey et al., 2011) have been classified as moderate quality, and values above 80% (Xin et al., 2020) or 90% (Dorsey et al., 2011) have been classified as high quality. The definition of motility also varies among researchers. In the present study we have defined motility as sperm with a VCL >10 μm/s (Xin et al., 2020; Shazada et al., 2024). Other studies have classified sperm as motile with a VCL >3 μm/s (Dzyuba et al., 2012; Shaliutina et al., 2013; Dzyuba et al., 2015; Barulin and Shumski, 2022), or as uncontrollably spinning in place (DiLauro et al., 1994), and some do not report their motility criteria (Park and Chapman, 2005; Aramli et al., 2013) making comparisons between studies difficult.

Longevity of sperm motility should also be considered when establishing motility criteria. For example, gamete contact time has a large effect on fertilization success (Butts et al., 2009), as sperm needs to be able to reach an egg. For sturgeon, their sperm can remain motile for up to several minutes and even hours in some cases (Ginzburg, 1972; Alavi et al., 2004; Gilroy and Litvak, 2019). Because of this relatively long period of motility, a VCL of both 3 μm/s or 10 μm/s may be sufficient to achieve fertilization. Therefore, defining minimum VCL for motility could significantly impact interpretation of sperm quality. It is clear that scientists working in this area of research need to specify motility criteria for comparisons within and between species.

While sperm in our study were still motile for up to 24 days, sperm after day 10 would be classified as poor quality in terms of fertilization capacity, as motility was less than 20% (Dorsey et al., 2011; Xin et al., 2020); after day 10, more than 80% of sperm had a VCL <10 μm/s (our motility definition).

Extenders tested in this experiment have been used to prolong sperm motility and kinematics in *Acipenser* sperm in previous studies. While all extenders used in these previous studies had similar pH and osmotic pressure there was still a noticeable difference in motility between extenders used (Table 4). Many of these studies found that the Park and Chapman extender had the longest lasting motility. This is likely because of the sugars used, and the phosphate content (Dorsey et al., 2013). Dorsey et al. (2013) speculated that sturgeon sperm cells possess glucose pores such as those found in sperm cells found in other species (Kim and Moley, 2008; Sancho et al., 2008), which demonstrates that glucose and phosphate are important for ATP stores and production, which are crucial for sperm motility. This would explain the difference in results we saw between HBSS and the Park and Chapman extender, as HBSS contains dextrose rather than glucose. This could also explain why sperm treated with the Park and Chapman extender in experiment 2 appear to decrease motility at a slower rate (prior to contamination) than the modified Tsvetkova extender, as the modified Tsvetkova extender contains sucrose rather than glucose.

**Table 4.** Results of short-term storage of *Acipenser* sperm from eight previous studies.

| Species | Motility<br>Decrease | Atmosphere | Extender | Days | Motility Definition | Author |
| --- | --- | --- | --- | --- | --- | --- |
| *A. persicus* | 100% | NO | N | 9 | Not mentioned | Aramli et al., 2013 |
| *A. baerii* | 100% | O (2 mg o2/L) | N | 5 | VCL>3 µm/s | Barulin and Shumski, 2022 |
| *A. baerii* | 100% | O (4 mg o2/L) | N | 6 | VCL>3 µm/s | Barulin and Shumski, 2022 |
| *A. baerii* | 100% | O (5.5 mg o2/L) | N | 8 | VCL>3 µm/s | Barulin and Shumski, 2022 |
| *A. baerii* | 100% | O (12 mg o2/L) | N | 11 | VCL>3 µm/s | Barulin and Shumski, 2022 |
| *A. oxyrinchus* | 60% | O | N | 17 | Cells uncontrollably spinning in place were classified as nonmotile | DiLauro et al., 1994 |
| *A. oxyrinchus* | 0% | O (96%) | P | 7 | Not mentioned | Dorsey et al., 2011 |
| *A. oxyrinchus* | 37.90% | NI (99%) | P | 7 | Not mentioned | Dorsey et al., 2011 |
| *A. oxyrinchus* | 0% | O (96%) | M | 7 | Not mentioned | Dorsey et al., 2011 |
| *A. oxyrinchus* | 95.20% | NI (99%) | M | 7 | Not mentioned | Dorsey et al., 2011 |
| *A. oxyrinchus* | 0% | O (96%) | N | 7 | Not mentioned | Dorsey et al., 2011 |
| *A. oxyrinchus* | 93.30% | NI (99%) | N | 7 | Not mentioned | Dorsey et al., 2011 |
| *A. oxyrinchus* | 40% * | O (100%) | PL | 21 | Not mentioned | Dorsey et al., 2011 |
| *A. oxyrinchus* | 50% * | O (100%) | PB | 21 | Not mentioned | Dorsey et al., 2011 |
| *A. oxyrinchus* | 70% * | O (100%) | N | 21 | Not mentioned | Dorsey et al., 2011 |
| *A. oxyrinchus* | 40% * | O (21%) | PL | 21 | Not mentioned | Dorsey et al., 2011 |
| *A. oxyrinchus* | 50% * | O (21%) | PB | 21 | Not mentioned | Dorsey et al., 2011 |
| *A. oxyrinchus* | 70% * | O (21%) | N | 21 | Not mentioned | Dorsey et al., 2011 |
| *A. ruthenus* | 30% * | NO | N | 1.5 | VCL>3 µm/s | Dzyuba et al., 2015 |
| *A. oxyrinchus desotoi* | 30.70% | NO | P | 21 | Not mentioned | Park and Chapman, 2005 |
| *A. brevirostrum* | 50% | NO | P | 28 | Not mentioned | Park and Chapman, 2005 |
| *A. oxyrinchus desotoi* | 100% | NO | N | 21 | Not mentioned | Park and Chapman, 2005 |
| *A. brevirostrum* | 100% | NO | N | 21 | Not mentioned | Park and Chapman, 2005 |
| *A. gueldenstaedtii* | 100% | NO | N | 9 | VCL>3 µm/s | Shaliutina et al., 2013 |
| *A. baerii* | 100% | NO | N | 9 | VCL>3 µm/s | Shaliutina et al., 2013 |
| *A. ruthenus* | 88.52% | NO | N | 4 | VCL>10 µm/s | Shazada et al., 2024 |
| *A. ruthenus* | 71.98% | NO | E17** | 6 | VCL>10 µm/s | Shazada et al., 2024 |
| *A. ruthenus* | 52.4% | NO | E18** | 6 | VCL>10 µm/s | Shazada et al., 2024 |
| *A. ruthenus* | 47.42% | NO | E19** | 6 | VCL>10 µm/s | Shazada et al., 2024 |
| *A. ruthenus* | 49.01% | NO | E20** | 6 | VCL>10 µm/s | Shazada et al., 2024 |
|  | *estimated from graphs<br>**see publication for extender recipe |  |  |  |  |  |
|  | N= No extender<br>P= Park and Chapman extender<br>M= Modified Tsvetkova extender<br>NO= No atmosphere treatment<br>O= Oxygen treatment<br>PL= Park and Chapman diluted after collection in lab<br>PB= Park and Chapman diluted immediately after collection |  |  |  |  |  |

In experiment 1 we did not increase oxygen content. For experiment 2 we added oxygen content as a factor, in addition to testing a different extender. Our results on the addition of oxygen were not conclusive, however we did see higher motility and swimming kinematic with sperm stored with oxygen after day 17 for all treatments (Figure 2b). In other studies, motility of *Acipenser* sperm stored in an oxygenated atmosphere was superior to that of other studies stored in an atmosphere with only air (DiLauro et al., 1994; Barulin and Shumski, 2022). However, the addition of oxygen can cause an increase of reactive oxygen species. The increase in both reactive oxygen species and heterotrophic protists is one of the most limiting factors of the storage life of sperm without cryopreservation (Barulin and Shumski, 2022), as was likely the case with experiment 2. While these protozoans did not appear to physically latch on or interact with the motile (or non-motile) sperm, it is likely that they would be competing for resources from the seminal plasma and extender mix (primarily the sugars used in the extenders). This would cause sperm longevity to decrease due to the lack of energy availability, as we observed in the samples treated with the Park and Chapman extender. It appears that while glucose is the preferred energy source for sperm stored in this experiment, it is likely also the preferred energy for the protozoans found in our experiment.

The most promising results from previous research were found by Park and Chapman (2005), who were able to maintain a long motility time for samples with a mixture of Shortnose sturgeon and Gulf of Mexico sturgeon sperm for 28 days without oxygen. This is likely because their study appears to be the only study on *Acipenser* sperm that used a shaker. Consistently shaking the stored milt in its tube allows for both oxygen and the fluid itself to be moved around within the tubes, therefore allowing sperm greater access to oxygen and resources from seminal fluid or added extenders. While their study was successful, it is important to consider that they did not examine sperm longevity separately by species. It may be that the presence of seminal fluid from Gulf of Mexico sturgeon enhanced longevity of the shortnose sturgeon sperm used in their experiment.

Even with the promising results from our study, future research should include the addition of a shaker to increase oxygen and particle movement within the stored seminal fluid, as well as testing antibiotics to lower the likelihood of contamination from bacteria.

## 5. Conclusion

Our findings lend insight into which extender is most effective during short-term storage for shortnose sturgeon sperm, how oxygen influences sperm longevity, and how these elements can contribute to developing a protocol for short-term storage of sperm in the field. This is crucial information to determine, because of the limited availability of sperm and the extreme risk of extinction for this vertebrate group. One of the main avenues to save sturgeons is artificial reproduction, which requires viable sperm that can be held until females can be stripped. Though cryopreservation is useful for banking sperm samples, it is impractical for field settings as the process is intricate and requires expensive and fragile equipment. Discovering methods to refine short-term storage protocols will allow sturgeon sperm to remain viable during transport for various applications such as assisted reproduction, reproductive research, and development of cryopreservation protocols. This work documents short-term storage of shortnose sturgeon sperm directly in the field using a mobile lab and demonstrates how this protocol can be further adjusted to be as practical and efficient as possible.

## Acknowledgments

The authors thank Danny Glassman, Alex Giroux, Naomi Meed, and Ahmad Hayat for their help with data collection.

## CRediT authorship contribution statement

Kenzie H. Melanson and Matthew K. Litvak: Conceptualization, Methodology, statistical analysis, writing, and editing.

